# *pixy*: Unbiased estimation of nucleotide diversity and divergence in the presence of missing data

**DOI:** 10.1101/2020.06.27.175091

**Authors:** Katharine L Korunes, Kieran Samuk

## Abstract

Population genetic analyses often use summary statistics to describe patterns of genetic variation and provide insight into evolutionary processes. Among the most fundamental of these summary statistics are *π* and *d*_XY_, which are used to describe genetic diversity within and between populations, respectively. Here, we address a widespread issue in *π* and *d*_XY_ calculation: systematic bias generated by missing data of various types. Many popular methods for calculating *π* and *d*_XY_ operate on data encoded in the Variant Call Format (VCF), which condenses genetic data by omitting invariant sites. When calculating *π* and *d*_XY_ using a VCF, it is often implicitly assumed that missing genotypes (including those at sites not represented in the VCF) are homozygous for the reference allele. Here, we show how this assumption can result in substantial downward bias in estimates of *π* and *d*_XY_ that is directly proportional to the amount of missing data. We discuss the pervasive nature and importance of this problem in population genetics, and introduce a user-friendly UNIX command line utility, *pixy*, that solves this problem via an algorithm that generates unbiased estimates of *π* and *d*_XY_ in the face of missing data. We compare *pixy* to existing methods using both simulated and empirical data, and show that *pixy* alone produces unbiased estimates of *π* and *d*_XY_ regardless of the form or amount of missing data. In sum, our software solves a long-standing problem in applied population genetics and highlights the importance of properly accounting for missing data in population genetic analyses.

## Introduction

Population geneticists often use summary statistics to describe patterns of genetic variation and to estimate population genetic parameters such as effective population size or mutation rate (Hartl et al. 1997; Gillespie 2004). The calculation of summary statistics is thus often the first step in a population genetic analysis, be it an exploratory study, a test of an evolutionary hypothesis, or the training of a machine-learning model (Hartl et al. 1997; Flagel et al. 2019; Hahn 2019). As such, accurate and unbiased algorithms for computing summary statistics are critical to the practice of population genetics.

Many summary statistics are based on the comparison of DNA sequences. Two important summary statistics in this class are *π*, the average number of nucleotide differences between genotypes drawn from the same population (Nei and Li 1979); and *d*_XY_, the average number of nucleotide differences between genotypes drawn from two different populations (Nei and Li 1979). These two summary statistics underlie a large variety of descriptive and inferential procedures in population genetics. For example, *π* is often used as an estimator of the central population genetic parameter (and is thus sometimes styled as). Similarly, *d*_XY_ is a key statistic for exploring patterns of divergence between populations, particularly in the context of divergence with gene flow (Noor and Bennett 2009; Cruickshank and Hahn 2014; Burri 2017).

### Calculation of π and d_XY_

For a single biallelic locus, *π* is usually calculated using one of three expressions shown in Equation 1, all of which are exactly equivalent:

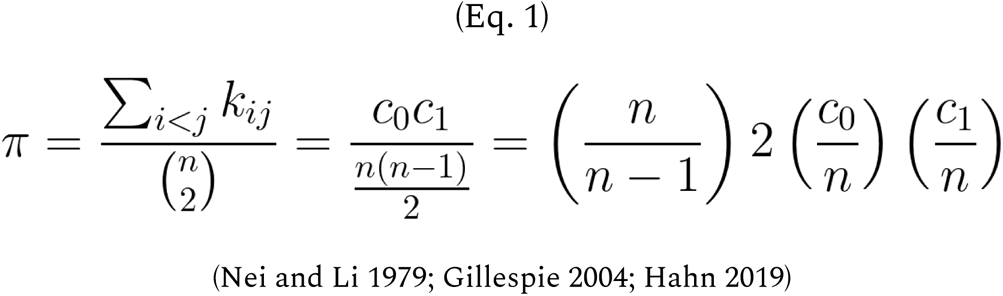

Where *k*_ij_ corresponds to the count of allelic differences between the *i*th and *j*th haploid genotypes, *n* is the number of samples, and c_0_ and *c*_*1*_ are the respective counts of the two alleles at the locus. Note that the last expression is simply the sample-size corrected expected heterozygosity (i.e. the “2pq” term in the Hardy-Weinberg equation).

*d*_XY_ is usually calculated using an all pairwise comparisons method (similar to the first expression in Eq. 1), with the only difference being that comparisons are only made between genotypes from different populations.

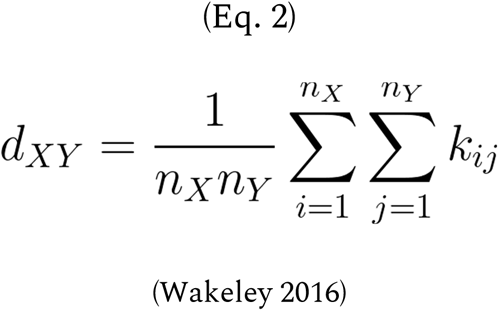

Where n_x_ and n_y_ correspond to the number of individuals in populations X and Y, and k_ij_ corresponds to the count of allelic differences between the *i*th (from population X) and *j*th (from population Y) haploid genotypes. There are also methods that use allele frequencies to approximate the result of Equation 2 (e.g. Smith and Kronforst 2013), but note that these ignore differences in sample size among sites.

Often, *π* and *d*_XY_ are computed for multi-locus genomic regions or in sliding windows. In order to standardize these statistics between sequences of different lengths, it is common to convert them to per-site estimates by dividing their raw value (computed for the whole sequence) by the total number of base pairs in the window (Hartl et al. 1997; Hahn 2019). However, two types of missing data can complicate this procedure (Figure 1). First, when genotype information at a site is missing in *all* samples, the sequence length must be adjusted accordingly downward. Second, when genotype information at a site is missing in *some* samples, the denominator of the raw value of *π* and *d*_XY_ (*n* in Equations 1 and 2) is *variable* across sites — a fact which must be accounted for in order to avoid the introduction of statistical bias in the final per site estimates (Nei and Roychoudhury 1974).

**Figure 1.**
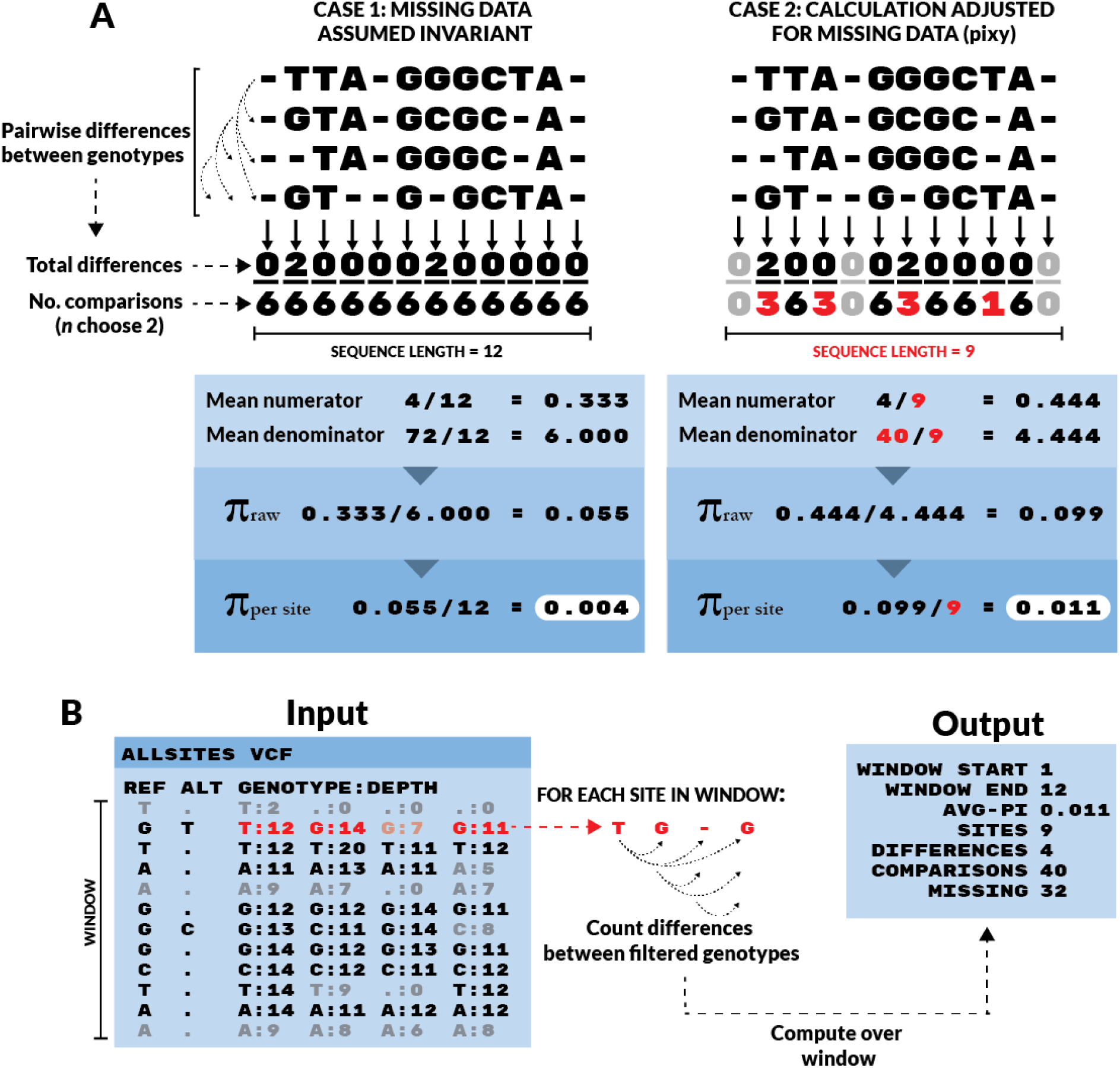
The logic and input/ouput of *pixy* demonstrated with a simple haploid example. (A) Comparison of two methods for computing *π* (or *d*_XY_) in the face of missing data. These methods follow the first expression of Equation 1 but differ in how they calculate the numerator and denominator. In Case 1, all missing data is assumed to be present but invariant. This results in a deflated estimate of *π*. In Case 2, missing data are simply omitted from the calculation, both in terms of the number of sites (the final denominator) and the component denominators for each site (the n choose 2 terms). This results in an unbiased estimate of *π*. (B) The adjusted *π* method (Case 2) as implemented for VCFs in *pixy*. Invariant sites are represented as sites with no ALT allele, and greyed-out sites are those that failed to pass a genotype filter requiring a minimum number of reads covering the site (Depth>=10 in this case).

Many population genetics texts introduce the calculation of *π* and *d*_XY_ using full sequence data (as seen in Figure 1A). When DNA data are represented in this way, the distinction between missing, variant (polymorphic), and invariant (monomorphic) sites is obvious and straightforward. However, modern population genomic data is rarely encoded in this way. In fact, one of the most common formats for encoding genomic data, the Variant Call Format (VCF), typically includes only sites that are genotyped as variant and does not usually explicitly distinguish invariant (but genotyped in the samples) sites from sites that are truly missing (Banks et al. 2011). The data summarized by VCFs can include both categories of missing data described above: sites that are entirely missing and genotypes that are missing within a site. The strategy of only reporting variant sites means that genotypes missing within a variant site are indicated, but sites that are entirely missing are omitted from the VCF and are thus indistinguishable from sites that were genotyped as invariant. This feature is usually intentional, as including information on millions of invariant sites massively increases file size and is superfluous for many analyses.

Unfortunately, information on both invariant and missing data is *not* superfluous for the calculation of *π* and *d*_XY_, and the absence of this information precludes the unbiased calculation of both statistics (Figure 1). This fact may be surprising to the reader, as many population genetics software tools provide methods of calculating *π* (and more rarely *d*_XY_) from variants-only VCFs. How then, do these tools distinguish missing from invariant sites, as is necessary for the unbiased calculation of per-site *π* and *d*_XY_? The answer to this question, as will be explored in depth here, is that the vast majority of existing tools make the simplifying assumption that missing sites are *present but invariant* (Case 1 in Figure 1). This assumption leads to downwardly biased estimates of *π* and *d*_XY_ in the presence of missing data. This approach is problematic for a variety of reasons: along with the general underestimation of *π* and *d*_XY_ it also creates a correlation between *π*/*d*_XY_ and “missingness”, which can itself covary with various features of the genome, e.g. TEs or structural variants (Carmena and González 1995; Kent et al. 2017).

The problematic nature of calculating *π* and *d*_XY_ in the presence of missing data is well known to practitioners of population genetics and is often overcome using a variety of ad-hoc methods. One common approach involves the creation of a VCF containing both invariant and variant sites (sometimes called an ‘all sites’ or ‘invariant sites’ VCF), from which information on truly missing sites can then be inferred (Burri 2017; Samuk et al. 2017; Irwin et al. 2018; Korunes et al. 2019). However, this approach has never been formalized as a general-purpose tool.

Here, we introduce *pixy*, a user-friendly command line utility for calculating *π* and *d*_XY_ from VCFs with invariant sites that correctly accounts for missing data. We compare the accuracy of *pixy*’s estimates of *π* and *d*_XY_ to those of existing methods using both simulated and empirical data. We show that *pixy* alone produces unbiased estimates of both statistics under a wide range of missing data conditions. More generally, we discuss the pervasive nature of missing data in population genetics, and use *pixy* to demonstrate the importance of accounting for it in the context of the calculation of *π* and *d*_XY_.

## New Approaches

*pixy* is a command-line tool, written in Python 3 and available on Github and via conda-forge for installation under Linux/OSX systems. The user supplies an ‘all sites’ VCF and a populations file listing the population(s) of interest and the associated sample names as listed in the VCF genotype columns. *pixy’s* documentation (https://pixy.readthedocs.io/en/latest/) provides guidance on VCF generation and filtering. *pixy* makes use of data structures provided by the Python module *scikit-allel* to efficiently handle invariant sites VCFs (Miles et al. 2019). The user can quickly compute π and *d*_XY_ over genomic windows of arbitrary size. The key difference between *pixy* and existing methods is the handling of missing data via dynamic adjusting of site-level denominators (which are propagated properly during windowed operations) and the adjustment of effective sequence length (Figure 1A). The output also includes all the raw information for all π and *d*_XY_ estimates (i.e., the component numerators and denominators for all computations). The name *pixy* is a play on the original parameter name π_XY_, which was used by (Nei and Li 1979) in place of *d*_XY_. All code is freely available on Github https://github.com/ksamuk/pixy, and detailed documentation is provided via readthedocs https://pixy.readthedocs.io/en/latest/. The software is also available for installation via Anaconda on the conda-forge channel https://anaconda.org/conda-forge/pixy.

## Results

### Validation of pixy results

To begin, we examined *pixy’s* accuracy as an estimator of π and *d*_XY_ by comparing *pixy’s* results to pre-existing methods and to theoretical expectations. We conducted 10,000 simulations to generate datasets in which all samples have observed genotypes at all sites (i.e., no missing data). Neutral simulations with a known effective population size and mutation rate allow us to compare *pixy’s* output to the simple theoretical expectation of E(π) = 4N_e_μ (Hartl et al. 1997; Wakeley 2016). To evaluate *d*_XY_, we split the simulated population into two random groups. In this case, 4N_e_μ is also the expected value of *d*_XY_ (this can be conceptualized as computing *d*_XY_ between two populations with a divergence time of zero, (Hahn 2019).

Using these simulated data, we compared the accuracy of *pixy’s* estimates of π and *d*_XY_ with several popular existing tools: VCFtools, PopGenome, and scikit-allel (Banks et al. 2011; Korneliussen et al. 2014; Pfeifer et al. 2014; Miles et al. 2019). These tools represent some of the most cited software packages for calculating π and *d*_XY_. Notably, the PopGenome manual acknowledges that π and *d*_XY_ estimates will be biased by missing data and includes a warning about computing estimates in the presence of missing data (Pfeifer et al. 2014). Nonetheless, users can and do use PopGenome to estimate π and *d*_XY_ in the presence of missing data, so we have chosen to include it here. Using each of these software packages, we computed π (and *d*_XY_ where available) for each of the simulated datasets. Using the resulting sampling distribution of π and *d*_XY_ from each estimation method, we compared the mean of each sampling distribution to the expected value of 4N_e_μ = 0.04 (Ne = 1×10^6^, μ =1×10^−8^). In the absence of missing data, all examined methods provide estimates of π and *d*_XY_ that closely match theoretical expectations (Figure 2 A-B; mean = 0.0398, standard error = 0.000189 for all sampling distributions). Estimates of π generated by *pixy* are identical to those of all other methods (R^2^ = 1.000, F=1.47×10^6^, df = 19998, p<2.2×10^−16^) and estimates of *d*_XY_ are nearly identical (R^2^ = 0.987, F=1.47×10^6^, df = 19998, p<2.2×10^−16^) (Figure 2 C-D). Overall, all compared methods have high accuracy when applied to complete datasets and provide nearly identical estimates when data are complete.

**Figure 2.**
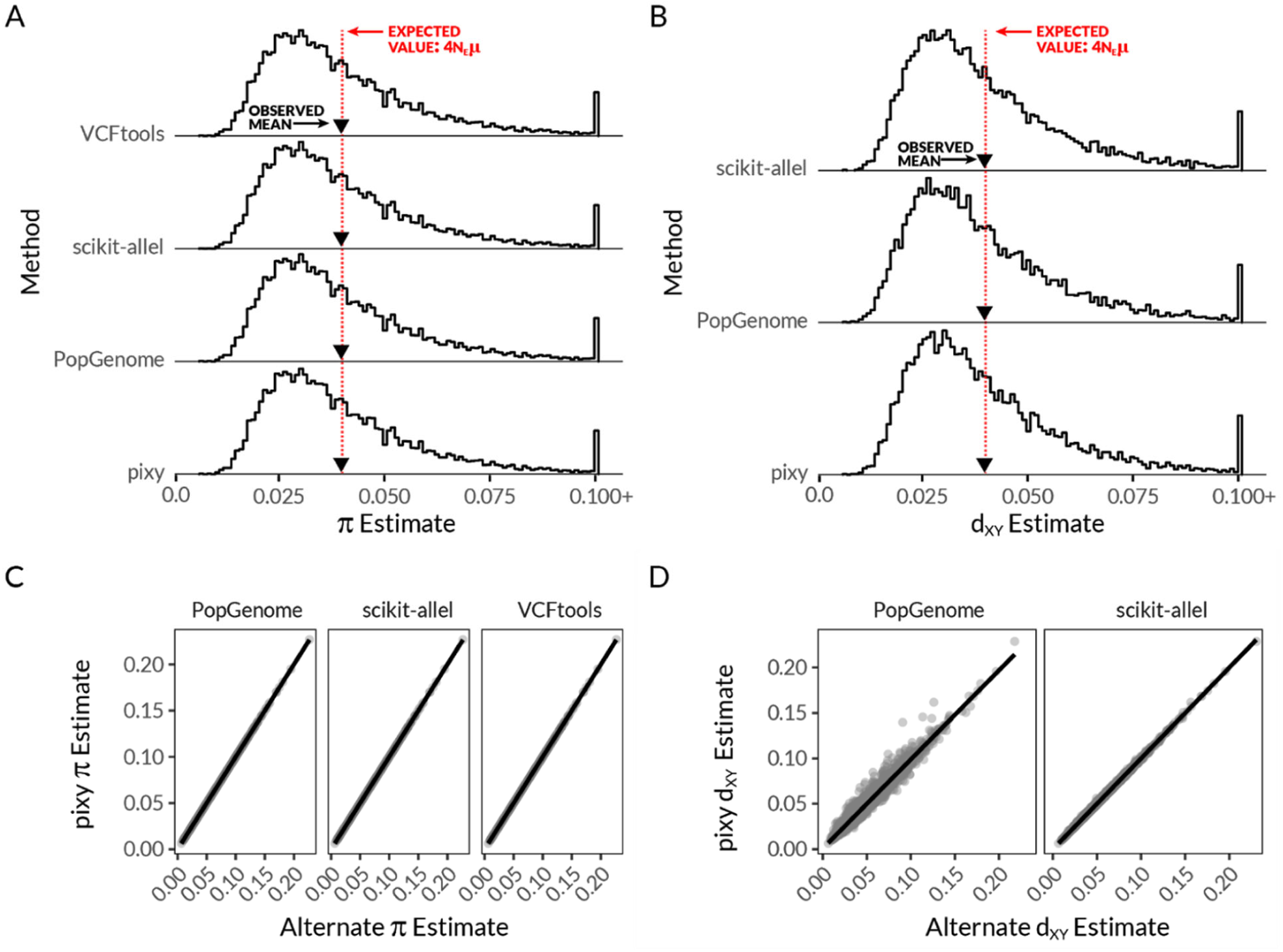
Comparison between *pixy* and existing methods in complete datasets. (A, B) The sampling distribution of π (A) and *d*_XY_ (B), as estimated from 10 000 simulated data sets using *pixy* and a variety of existing methods (see text for details). The red dotted line denotes the theoretical expectation for the mean of the sampling distribution, 4N_e_μ = 0.04 (which is the same for π and *d*_XY_ in this particular case). The observed means of the sampling distributions are marked with inverted triangles. For clarity, estimates of π and *d*_XY_ above 0.100 are aggregated in the last bin (“0.100+”). (C,D) direct comparisons between pixy’s estimates of π (C) and *d*_XY_ (D) and those from existing methods.

### pixy is unbiased in the presence of missing data

We next examined *pixy’s* accuracy in the presence of missing data. For a random set of 100 of the previously simulated datasets, we created subsets with varying quantities of missing data. This means that for each dataset with missing data, we had a corresponding “parent” dataset with complete data. Again, we computed π and *d*_XY_ for each dataset with the popular existing programs VCFtools, PopGenome, and scikit-allel.

In order to better visualize the effects of missing data, we scaled estimates of π and *d*_XY_ for each dataset with missing data by dividing by the estimate obtained from the parent dataset with no missing data. This normalizes any initial differences in π among datasets due to sampling variance. After this normalization, we observed that *pixy*’s π and *d*_XY_ estimates remain unbiased in the face of missing data (Figure 3, left column). As the proportion of missing data increases, the variance in estimates of π and *d*_XY_ increases. This spread in estimates across both sides of the y=1 line in Figure 3 increases as a function of missing data. Note, however, that the mean (expected) values of π and *d*_XY_ for *pixy* do not exhibit any significant trend (flat red lines, *pixy* panels, Figure 3, linear model slope does not differ significantly from 0 for sites or for genotypes, p > 0.2, Table S2). This is the expected behavior of an unbiased summary statistic in the face of missing data. In contrast, the three other methods all display a downward bias in their estimates of π and *d*_XY_ that increases as a function of the proportion of missing sites or genotypes (non-pixy panels, Figure 3; all slopes significantly negative for missing sites and genotypes, all p<2.2×10^−16^, Table S2). The effect of this bias was strongest for the case of completely missing sites, whereas missing genotypes (sites with missing genotypes for some samples) only begin to display strong bias around 80% missing data for most methods (Figure 3). The notable exception to this was VCFtools, which displayed a sharper increase in bias for ‘missing genotypes’ than ‘missing sites’ (Figure 3).

**Figure 3.**
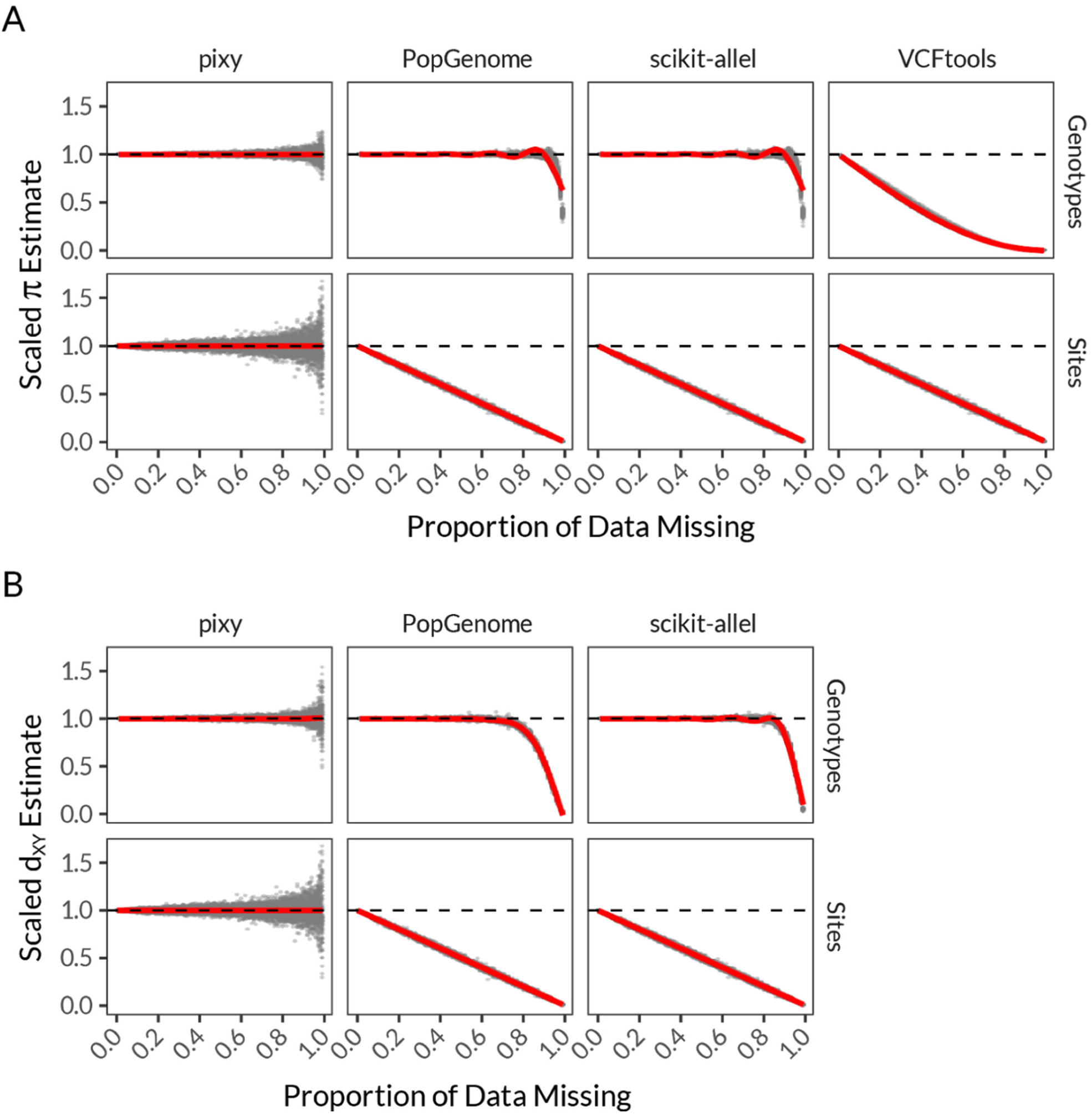
Comparison between *pixy* and existing methods in the presence of missing data. π (A) and *d*_XY_ (B) are shown as scaled estimates (each estimate is scaled by dividing by the estimate obtained from the parent dataset with no missing data). Perfect congruence between estimates in the presence and absence of missing data is shown with the dotted line at y=1. Estimates were obtained from datasets with varying proportions of missing genotypes (top row, A and B) and sites (bottom row, A and B).

### Analysis of empirical data: Anopheles gambiae

Finally, we applied *pixy* to an empirical dataset: deep sequencing of *Anopheles gambiae* provided by the Ag1000 Genomes Consortium. We used *pixy* to generate windowed estimates of π on the X chromosome for a sample (n=18) of the *A. gambiae* Burkina Faso (BFS) population, and we compared these estimates to those generated by popular pre-existing methods. We also examined *d*_XY_ between the 18 BFS samples and 18 additional samples from the KES (Kenya) population (Table S2). In addition to the three previously explored programs (VCFtools, PopGenome, and scikit-allel), we included estimates of π and *d*_XY_ from the software ANGSD (Korneliussen et al. 2014). ANGSD relies on genotype likelihoods calculated using the reads covering a position, making it incompatible with our simulated data but equipped to handle empirical sequencing data.

All four methods yielded estimates of π that were correlated with *pixy’s* estimates (Figure 4, R^2^ = 0.68 for VCFtools, 0.82 for ANGSD, and 0.79 for both PopGenome and scikit-allel). However, the previously identified biases caused by missing data appeared to result in substantial differences in estimates of π in many cases (Figure 4). In general, the compared methods tend to underestimate π, with the exception being ANGSD. This downward bias is seen as the grouping of estimates above the y=x line in Figure 4 A, C-D. As expected, the magnitude of this bias was closely correlated with the proportion of missing data (Figure 4, Figure S2). For the relatively complete regions of our subset of the Ag1000g dataset, the apparent underestimation of π was low (around −5%), but rapidly increased in cases of even moderate missingness (e.g. as much as −95% in cases of just 25% missing data, Figure S2). As expected, the same pattern of bias was also apparent for estimates of *d*_XY_ (Figure S1, Figure S3).

**Figure 4.**
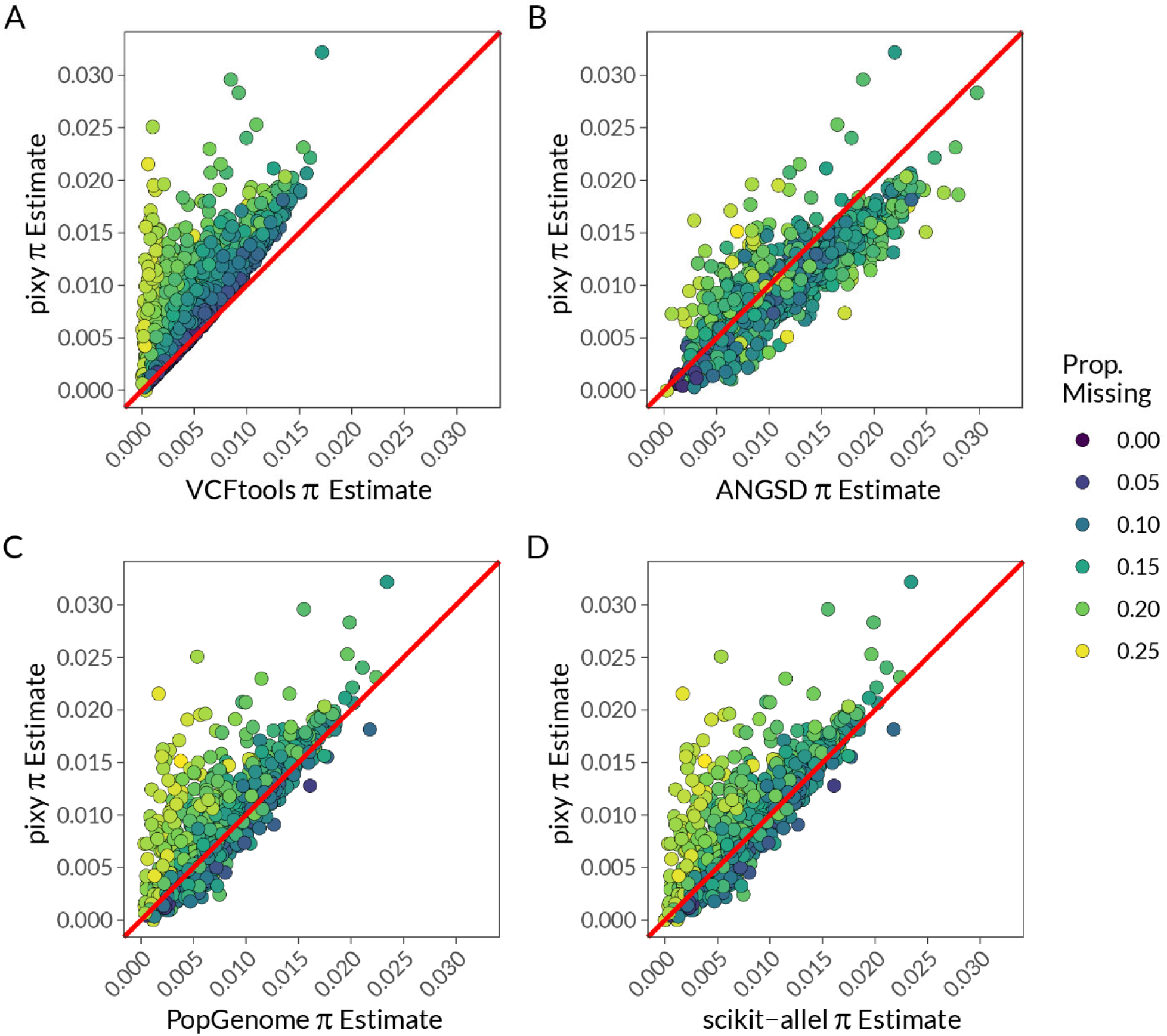
Comparisons of estimates of π from whole genome data derived from 18 *Anopheles gambiae* individuals from the Ag1000G Burkina Faso (BFS) population. Each panel depicts the estimates of π for the X chromosome performed using *pixy* (y-axis) and four other methods (x-axis, A-D). Points are colored according to the proportion of missing data (of any type) calculated by pixy. The 1:1 line is shown in red.

## Methods

### Simulated data: coalescent simulations via msprime

To provide ground-truthed datasets for evaluating the performance of *pixy*, we simulated sequence data using the coalescent simulator msprime (Kelleher et al. 2016). We created 10 000 simulated datasets, each with the following parameters: Effective population size = 1×10^6^, mutation rate =1×10^−8^, sample size = 100, number of sites = 10 000. We converted these to VCFs with invariant sites using a custom script (see code supplement). These datasets represent the case of “no missing data”. To explore the effects of different types of missing data, we randomly selected 100 of these original datasets as “parent” VCFs and then used these to simulate variable proportions of missing data (ranging from 0.0 to 0.99, in steps of 0.01). To simulate missing sites, we randomly dropped rows from each parent VCF. To simulate missing genotypes, we randomly converted a fixed proportion of genotypes in each parent VCF at every site to “./.” (missing). This resulted in a total of 30,000 VCFs of simulated data (10 000 original VCFs, and 10 000 “missing sites” VCFs and 10 000 “missing genotypes” VCFs).

### Empirical data: *Anopheles gambiae* whole genome data

To evaluate the performance of pixy in a realistic use case, we obtained short-read whole-genome sequencing data from two populations of *Anopheles gambiae* sequenced by the Ag1000G (*Anopheles gambiae* 1000 Genomes) Consortium (MalariaGEN 2016). We selected 18 individuals each from two populations: BFS (Burkina Faso) and KES (Kenya). Sample accessions and sequencing details are provided in Table S1. All sequences were aligned to the *Anopheles gambiae AgamP4*.*12* reference genome (Giraldo-Calderón et al. 2015) using BWA-0.7.5a (Li and Durbin 2009), after using Picard to mark adapters and duplicates (Broad Institute 2019). Variants were called using GATK version 4.1.1.0, using the “-all-sites” flag of GenotypeGVCFs to generate ‘all sites’ VCFs for input into pixy (McKenna et al. 2010; Van der Auwera et al. 2013)

### Comparison to existing methods using both simulated and empirical data

We compared the accuracy of *pixy’s* estimates of π and *d*_XY_ with several popular existing tools: VCFtools, ANGSD, PopGenome, and scikit-allel (Banks et al. 2011; Korneliussen et al. 2014; Pfeifer et al. 2014; Miles et al. 2019). We computed π using *pixy*, VCFtools, PopGenome, scikit-allel, and ANGSD. Note that ANGSD was only applied to the empirical data, since its diversity functions are not designed to work with VCFs. We computed *d*_XY_ using *pixy*, PopGenome, scikit-allel, and the ANGSD companion script “calcDxy” (Penalba 2016, https://github.com/mfumagalli/ngsPopGen/blob/master/scripts/calcDxy.R). For VCFtools, we used the ‘--window-pi’ method to estimate windowed π. For scikit-allel, we used the allel.sequence_diversity and allel.sequence_divergence functions to estimate windowed π and *d*_XY_, respectively. For PopGenome, we used the nuc.diversity.within and nuc.diversity.between functions, following the recommendations in the manual. We stress that the PopGenome manual explicitly warns that computing π and *d*_XY_ in the presence of missing data will result in biased estimates (Pfeifer et al. 2014). We have chosen to include it here because PopGenome is commonly used to estimate π and *d*_XY_ in spite of this warning.

We first used our simulated datasets to examine *pixy’s* accuracy in comparison to these existing methods. To obtain two simulated populations for evaluating *d*_XY_, we split the 100 simulated samples into two random groups. To standardize sample sizes between π and *d*_XY_ estimates, we computed π using the first half of the simulated individuals (n=50), and *d*_XY_ by designating the first half of the individuals as drawn from “Population 1” and the second half as drawn from “Population 2” (each with n=50 individuals). We computed π and *d*_XY_ in 10kb windows in each of the VCFs with variable missing data. *pixy* was run using default settings, and each pre-existing method was applied using the functions described above (see code supplement).

We then compared the accuracy of each method using the empirical *Anopheles gambiae* data. To do this, we first applied a basic genotype-level hard filter (DP >=10, GQ>=40|RGQ>=40) to the invariant sites VCF produced by GATK. This VCF was the input file for all methods apart from ANGSD (see below). We then computed π (*pixy*, VCFtools, ANGSD, PopGenome, scikit-allel) and *d*_XY_ (*pixy*, PopGenome, scikit-allel, ANGSD) in 10kb windows. We computed π separately for the BFS and KES populations. For ANGSD, the BAM files generated from the *Anopheles* BFS and KES populations were used as input, resulting in estimates of both π (ANGSD’s “pairwise theta”) and *d*_XY_ (obtained via a companion script: calcDxy – Penalba 2016, https://github.com/mfumagalli/ngsPopGen/blob/master/scripts/calcDxy.R). In the case of *π*, we explicitly divided the raw estimates of pairwise theta by the number of sites (nSites) provided by ANGSD, and not the window size (10000).

## Discussion

Modern population genomic analyses frequently rely on *π* and *d*_XY_ as measures of diversity and divergence, but these summary statistics are deceptively difficult to accurately calculate (Hartl et al. 1997; Gillespie 2004). Specifically, the correct handling of missing and invariant sites presents a common pitfall in the calculation of *π* and *d*_XY_. This challenge stems in part from the way genetic variation data is commonly encoded. The widely used Variant Call Format typically condenses data down to only variant sites and does not maintain information about which sites had insufficient data for genotyping and which sites were genotyped as invariant (Banks et al. 2011). If *π* and *d*_XY_ are calculated under the assumption that missing data are invariant, then the resulting estimates of *π* and *d*_XY_ are likely to be downwardly biased in many cases.

We observe this downward bias in our application of several popular tools using both simulated and empirical datasets. While many population geneticists recognize that such tools must be applied with caution, the lack of formalized best practices and available software for calculating *π* and *d*_XY_ in the presence of missing data leads to inconsistent approaches across studies. It also places the onus on the user to devise ad-hoc methods to handle missing data when using common software. *pixy* provides a user-friendly command line utility for estimating *π* and *d*_XY_ in a manner unbiased by the presence of missing data. We leverage a common strategy for distinguishing missing and invariant sites by (1) making use of VCFs including invariant sites and (2) employing algorithms that explicitly account for missing data. More generally, our comparison of *pixy* to existing tools demonstrates the consequences of failing to handle missing data properly and underscores the potential pervasiveness of this problem in population genetics.

In addition to generally underestimating *π* and *d*_XY_, failing to properly handle missing data can also create a correlation between *π*/*d*_XY_ and “missingness”, or the proportion of missing genotypes. This relationship is noteworthy since missingness is often tied to various features of the genome or of the data itself. For example, genomic features such as transposable elements or structural variation can cause variable assembly and mapping quality (O’Leary et al. 2018). Relatedly, individuals within a sample often have variable missingness (e.g., due to sample quality variation), which can generate false differences in *π* and *d*_XY_ if these vary systematically among biological units (e.g. between populations). Differences in genomic library preparation technique can also affect missingness, for example genome complexity reduction techniques (e.g., RAD-Seq or GBS) may present particularly variable and high levels of missingness relative to high-coverage whole-genome sequencing (Elshire et al. 2011; Lowry 2017). However, as seen with our case study of *Anopheles gambiae* data, even ∼30X coverage whole-genome sequencing can exhibit significant downward bias in *π* estimation when missing data is not taken into account (e.g., Figure S2). This is concerning, as this relatively high-quality, well-curated data set is likely close to a “best case” scenario for missingness, and most datasets will likely fare much worse. Given these considerations, we argue that best practices for calculating *π* and *d*_XY_ should always explicitly account for missing data.

One notable exception to the patterns we identified here was the likelihood-based method ANGSD (Korneliussen et al. 2014). ANGSD did not appear to display a systematic underestimation of *π* or *d*_XY_ in the face of missing data. However, the estimates produced by ANGSD are not fully congruent with *pixy*, and ANGSD appears to potentially systematically *overestimate π* or *d*_XY_ in some scenarios (Figure S2). While it is outside the scope of our current effort to systematically explore how ANGSD reacts to missing data, we note that it employs a rather different approach to analysis by working with allele frequencies rather than directly using genotypes. It is also important to note that the calculation of *d*_XY_ using ANGSD required post-processing using a third party script, as well as further processing using a custom script (written by the authors) to average over windows (see code supplement). The lack of a single validated protocol for calculating *d*_XY_ (or even *π*) using ANGSD suggests there may be a great deal of interstudy variation in estimates produced with ANGSD. It also differs from the other software used here in that ANGSD was not designed to analyze VCFs (though recent versions do input and output VCFs with some limitations), and thus may be more difficult to apply to many datasets.

While the unbiased methods provided by *pixy* are an important resource for facilitating *π* and *d*_XY_ calculation, much work remains to correct systemic issues in estimating diversity and divergence. Future studies are needed to address how missing data may affect the wide variety of other population summary statistics and tests (e.g. Wong et al. 2019). Another important area of future work is the development of file formats that efficiently store genetic data while maintaining the ability to distinguish well-supported invariant sites from sites which have insufficient information to determine whether they are truly invariant. As the field of population genetics advances, we hope that articulating this issue will provide groundwork for handling missing data as new file formats arise and new tools are developed.

## Acknowledgements

We thank the Ag1000G (*Anopheles gambiae* 1000 Genomes) Consortium for making their dataset publicly available and welcoming its use for testing purposes. Jerome Kelleher assisted in adapting msprime to create ‘all sites’ VCFs. Sarah Marion provided helpful discussions and the full derivation of Equation 1. David Peede and Brian Myers identified important file handling bugs in pre-releases of *pixy*. Mohamed Noor provided key resources and mentorship for both authors throughout the project, and KLK was supported by mentorship and resources provided by Amy Goldberg. The Noor Lab and Ross-Ibarra Lab provided helpful comments on early versions of this manuscript. This work was supported by: the Natural Sciences and Engineering Research Council of Canada via a Postdoctoral Fellowship awarded to KS; the National Science Foundation grant DEB-1754439 awarded to M. Noor; and the National Institute of General Medical Sciences at the National Institutes of Health grant 1R35GM133481-01 awarded to Amy Goldberg.

## Supplemental Material

**Figure S1.**
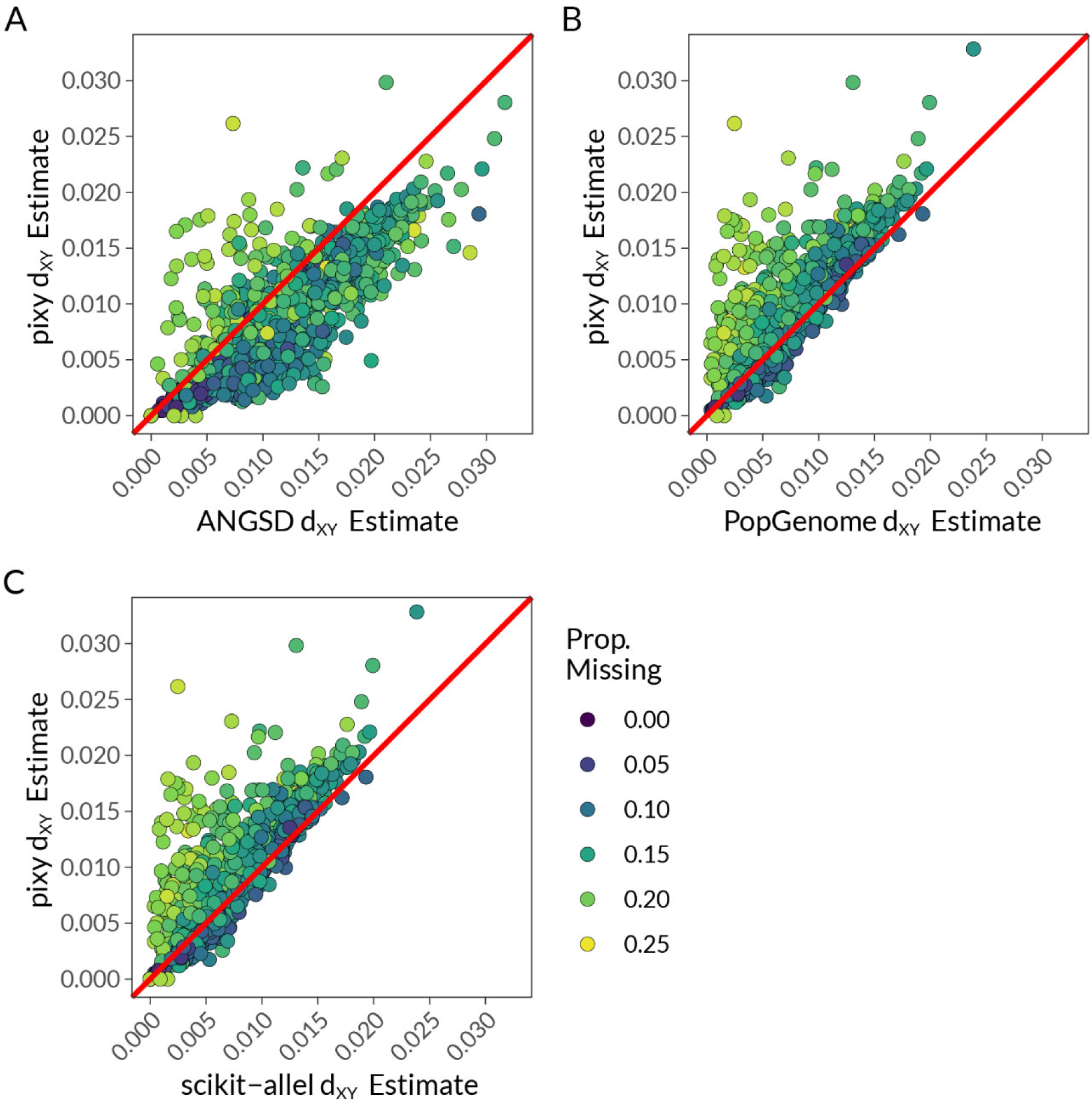
Comparisons of estimates of *d*_XY_ from whole genome data derived from 18 *Anopheles gambiae* individuals from the Ag1000G Burkina Faso (BFS) and the Kenya (KES) populations. Each panel depicts the estimates of *d*_XY_ for the X chromosome performed using pixy (y-axis) and four other methods (x-axis, A-D). Points are colored according to the proportion of missing data (of any type) calculated by pixy. The 1:1 line is shown in red.

**Figure S2.**
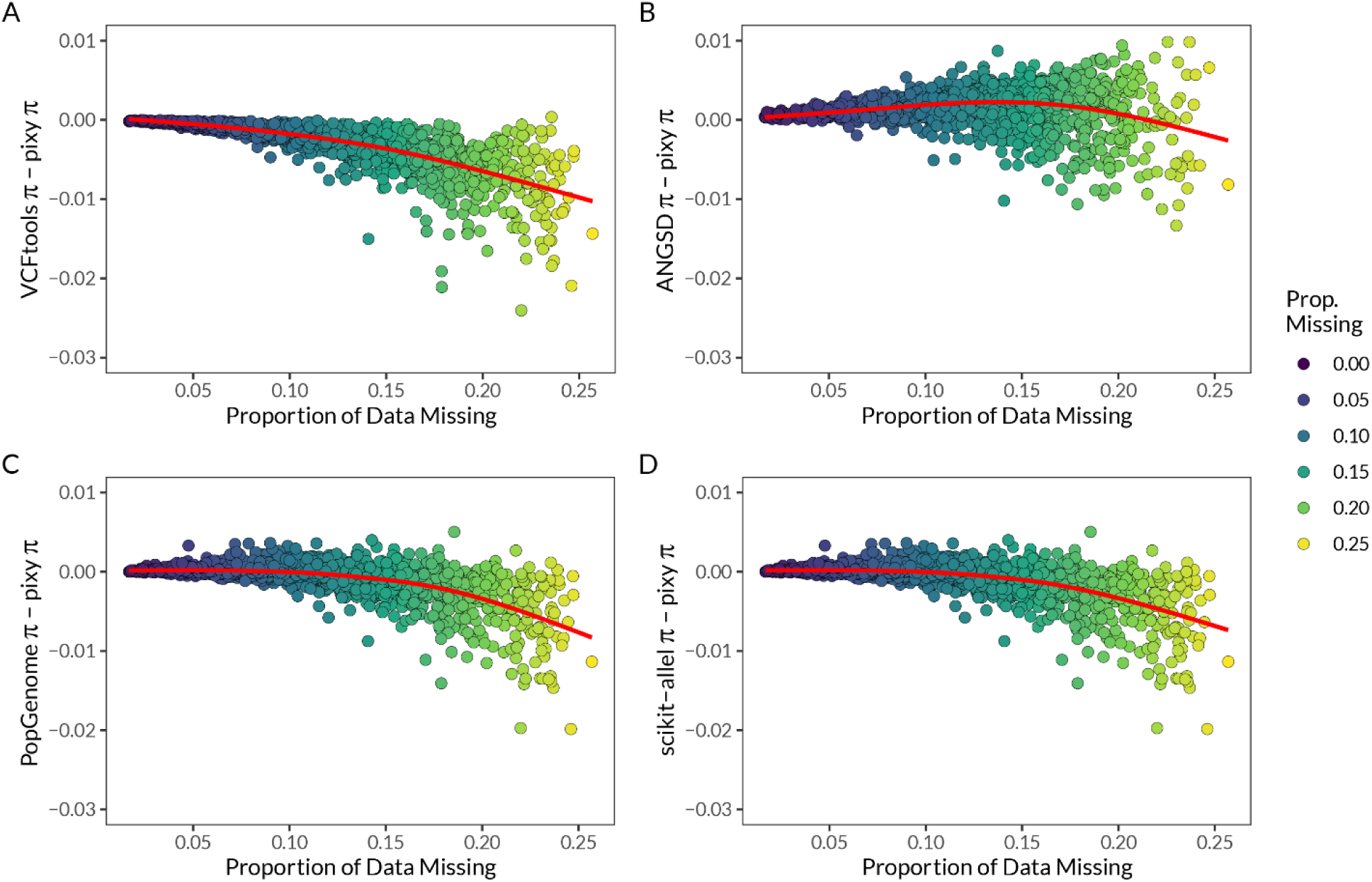
The difference in estimates of π between pixy and four existing methods as a function of missing data. All estimates derived from the X chromosome of 18 *Anopheles gambiae* individuals from the Ag1000G Burkina Faso (BFS) population (the same data are plotted in Figure 4). The y-axis is the difference in π estimates (method minus pixy), i.e. negative values suggest underestimation, assuming pixy is unbiased. Points are colored according to the proportion of missing data (of any type) calculated by pixy. Red lines are LOESS smoothed means.

**Figure S3.**
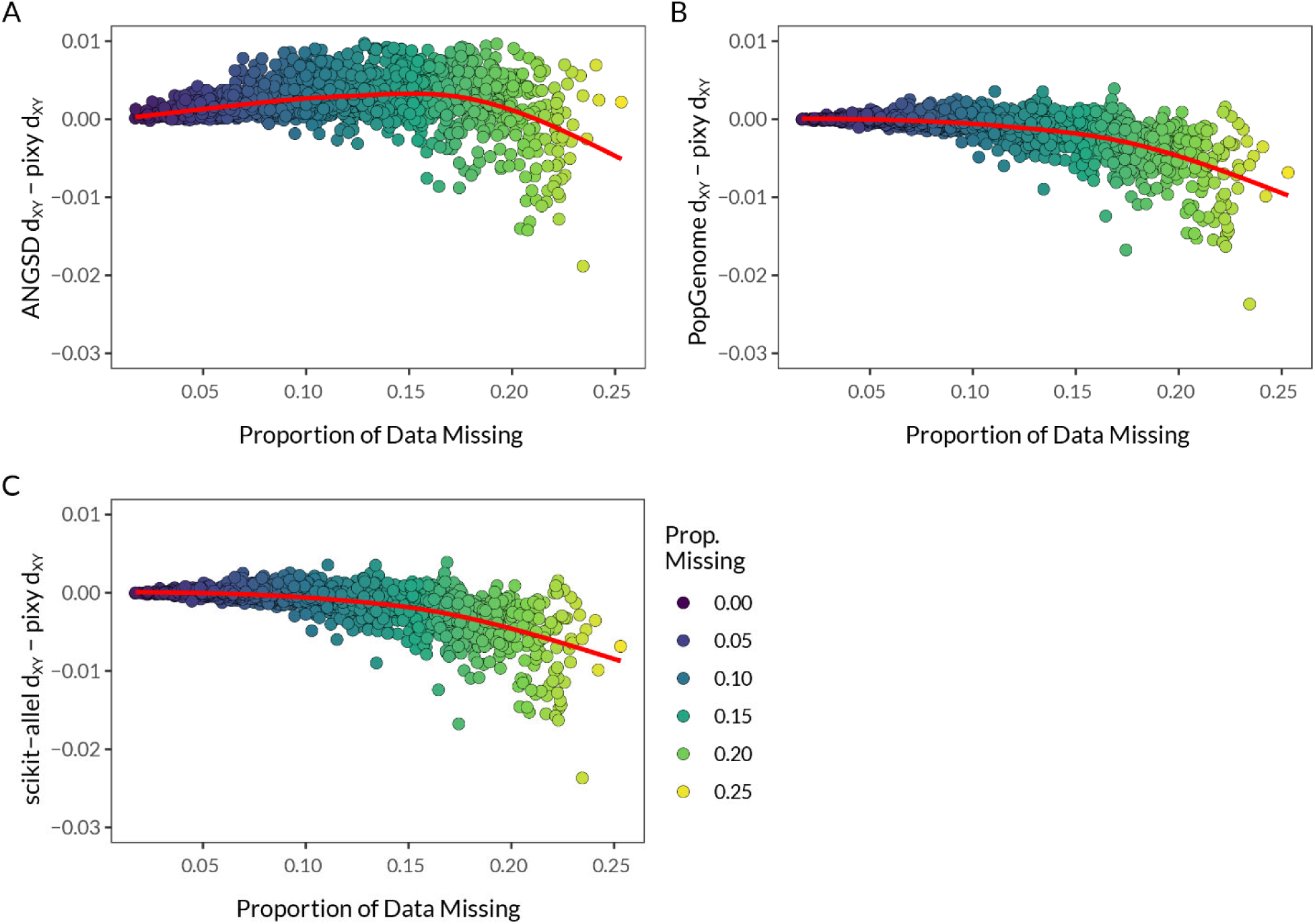
The difference in estimates of *d*_XY_ between pixy and four existing methods as a function of missing data. All estimates derived from the X chromosome of 18 *Anopheles gambiae* individuals from the Ag1000G Burkina Faso (BFS) and the Kenya (KES) populations (the same data are plotted in Figure S1). The y-axis is the difference in *d*_XY_ estimates (method minus pixy), i.e. negative values suggest underestimation, assuming pixy is unbiased. Points are colored according to the proportion of missing data (of any type) calculated by pixy. Red lines are LOESS smoothed means.

## Supplemental Table Captions

**Table S1** | Sample data and accessessions for *Anopheles gambiae* genomic data sourced from the Ag1000G (*Anopheles gambiae* 1000 Genomes) Consortium AR3 data release. This metadata is a subset of the full metadata set provided at https://www.malariagen.net/apps/ag1000g/phase1-AR3/index.html#start.

**Table S2** | Regression coefficients of linear models regressing four different methods of estimates of *π* and *d*_XY_ against the proportion of missing data (as estimated by pixy). Each row is shows the regression coefficient, standard error, T-test statistic vs. a null hypothesis of 0, and p-value of the resulting test for a single combination of statistic (pi or dxy), software method, and missing data type (genotypes = partially missing, sites = fully missing).

